# Influence of nanobody binding on fluorescence emission, mobility and organization of GFP-tagged proteins

**DOI:** 10.1101/2020.06.11.146274

**Authors:** Falk Schneider, Christian Eggeling, Erdinc Sezgin

## Abstract

Advanced fluorescence microscopy studies require specific and monovalent molecular labelling with bright and photostable fluorophores. This necessity led to the widespread use of fluorescently labelled nanobodies against commonly employed fluorescent proteins. However, very little is known how these nanobodies influence their target molecules. Here, we observed clear changes of the fluorescence properties, mobility and organisation of green fluorescent protein (GFP) tagged proteins after labelling with an anti-GFP nanobody. Intriguingly, we did not observe any co-diffusion of fluorescently-labelled nanobodies with the GFP-labelled proteins. Our results suggest significant binding of the nanobodies to a non-emissive, oligomerized form of the fluorescent proteins, promoting disassembly into more monomeric forms after binding. Our findings show that great care must be taken when using nanobodies for studying dynamic and quantitative protein organisation.

## Introduction

Labelling a protein of interest with an antibody is a well-established procedure in molecular biology. Rather large size and multivalence of antibodies, however, limit their application as labelling agents in imaging approaches. Over the past years, the popularity of antigen-binding fragments of antibodies (Fabs) and single-chain nanobodies derived from camelids or shark antibodies grew vastly (Beghein and Gettemans, 2017; Carrington et al., 2019; Leslie, 2018). Both types of molecules are much smaller than full-length antibodies, yet possess similar binding properties to their target proteins (Harmsen and De Haard, 2007; Sahl et al., 2017). Moreover, they only have a single binding site which prevents cross-linking and artificial clustering (Pereira et al., 2019; Sograte-Idrissi et al., 2020; Stanly et al., 2016). Additionally, the stoichiometric labelling of full length antibodies is challenging, whereas fluorescent labelling of a nanobody with 1:1 (nanobody:dye) ratio is regularly achieved (Grußmayer et al., 2014). Nanobodies have successfully been raised against various target molecules and used in microscopy (Pleiner et al., 2015, 2018). Some examples for nanobody epitopes include histones (Jullien et al., 2016), viral protein (Cao et al., 2019), artificial peptides (Braun et al., 2016), clathrin coat components (Traub, 2019), vimentin (Maier et al., 2015) and many more (Aguilar et al., 2019; Mikhaylova et al., 2015). Interestingly, a study using nanobodies targeting synaptic proteins and making use of the nanobodies’ smaller size and better penetration capabilities suggested a new pool of synaptic vesicles (Maidorn et al., 2019). The production methods and costs of generating a novel nanobody are comparable to the ones for a standard monoclonal antibody, however, the nanobody can subsequently be produced and harvested from bacteria, yeast or mammalian cell culture and even recombinantly tagged (Arbabi Ghahroudi et al., 1997; Beghein and Gettemans, 2017; Pleiner et al., 2018).

The use of nanobodies in microscopy was fuelled by the development of a green fluorescent protein (GFP) binding nanobody (Ries et al., 2012). GFP or its derivatives (like the enhanced GFP, EGFP) are attractive targets for super-resolution microscopy as they can be considered the biologist’s favourite tag, and a GFP-tagged version of a protein of interest is routinely cloned. However, compared to organic dyes, the brightness and photostablity of GFP and its variants is usually worse, limiting its use in some applications. Here, the use of anti-GFP nanobodies labelled with, for example, an organic dye with desired chemical or photophysical properties paved the way for a variety of applications (Beghein and Gettemans, 2017; Buser et al., 2018; Fabricius et al., 2018; Farrants et al., 2020; Ries et al., 2012; Sahl et al., 2017) and the development of nanobodies against other fluorescent proteins (FPs) (Platonova et al., 2015). This, in turn, allowed, for example, for multi-colour super-resolution imaging with nanobodies (Sograte-Idrissi et al., 2019).

The binding of the anti-GFP nanobody to GFP has been characterized (Kirchhofer et al., 2010; Klamecka et al., 2015; Della Pia and Martinez, 2015) and it has already been noted that the binding of a nanobody to a fluorescent protein can change the photophysical properties of GFP such as fluorescence brightness, depending on the binding site (Kirchhofer et al., 2010). This influence has been exploited for in vivo studies (Llama Tags in fruit fly embryo (Bothma et al., 2018)). General fluorescence properties of GFP have been studied in depth (Conyard et al., 2011; Jung et al., 2005), and its fluorescence brightness and lifetime, as well as excited- and dark-state populations have been shown to depend on environmental characteristics such as solvent properties (e.g. pH, viscosity), illumination intensity, and wavelength (Ghosh et al., 2017; Jung et al., 2005; Lippincott-Schwartz and Patterson, 2009; Niwa et al., 1996; Tsien, 1998).

Influences of the nanobody on the functionality of the FP-tagged protein have been indicated before (Küey et al., 2019), and we here present new insights by investigating effects of nanobody binding on the fluorescence emission, organization and mobility of GFP-tagged proteins. Specifically, we used commercially available unlabelled and fluorescently labelled GFP-binding nanobodies (Nb) in combination with fluorescence imaging and spectroscopic tools such as fluorescence correlation spectroscopy (FCS) for GFP and EGFP in solution, attached to synthetic membranes, and expressed on the surface of live cells as (E)GFP-tagged GPI-anchored proteins. Our data suggests that the anti-GFP Nb binds a dark oligomeric form of GFP and promotes reorganization by releasing bright monomers.

## Results and Discussion

### Nanobody binding in solution

We first investigated the basic fluorescence properties of GFP and EGFP before and after addition of unlabelled nanobody (Nb) in solution. Specifically, using a fluorescence spectrometer we investigated changes in total fluorescence intensity and fluorescence spectra for recombinant his-tagged (E)GFP in PBS (pH 7.4, room temperature). Figure 1 shows the respective excitation and emission spectra. As reported before (Tsien, 1998), we found two excitation peaks at around 395 nm and 480 nm for GFP corresponding to the neutral and deprotonated (anionic) state of the fluorochrome, respectively (Chattoraj et al., 1996; Chudakov and Lukyanov, 2003), and one excitation peak at around 480 nm for EGFP (Figure 1a,b). The anionic state is usually considered for fluorescence microscopy experiments, using a standard 488 nm laser line for excitation. Due to the requirement for UV excitation, the neutral form of GFP is less used. Interestingly and already previously indicated (Kirchhofer et al., 2010), Nb binding promotes anionic state absorption, revealed by a ≈ 2-fold reduction of the excitation peak at 390 nm and a corresponding ≈ 3-fold increase at 480 nm for GFP (Figure 1a). Similarly but less pronounced, the excitation peak at around 480 nm also increased by 25% for EGFP upon interaction with the Nb (Figure 1b). The Nb binding did not induce any shifts in the emission spectra of GFP or EGFP (excited with 488 nm, Figure 1c,d), also not under UV excitation (405 nm excitation, Figure 1e,f), i.e. peak positions of the spectra remained the same with inefficient excitation of EGFP at 405 nm. Overall, in solution GFP and EGFP experience a ≈ 3.5– and ≈ 1.5–fold increase in total integrated fluorescence emission (510 nm to 600 nm) induced by the Nb binding, apparently mainly due to the increase in absorption at around 480 nm.

**Figure 1:**
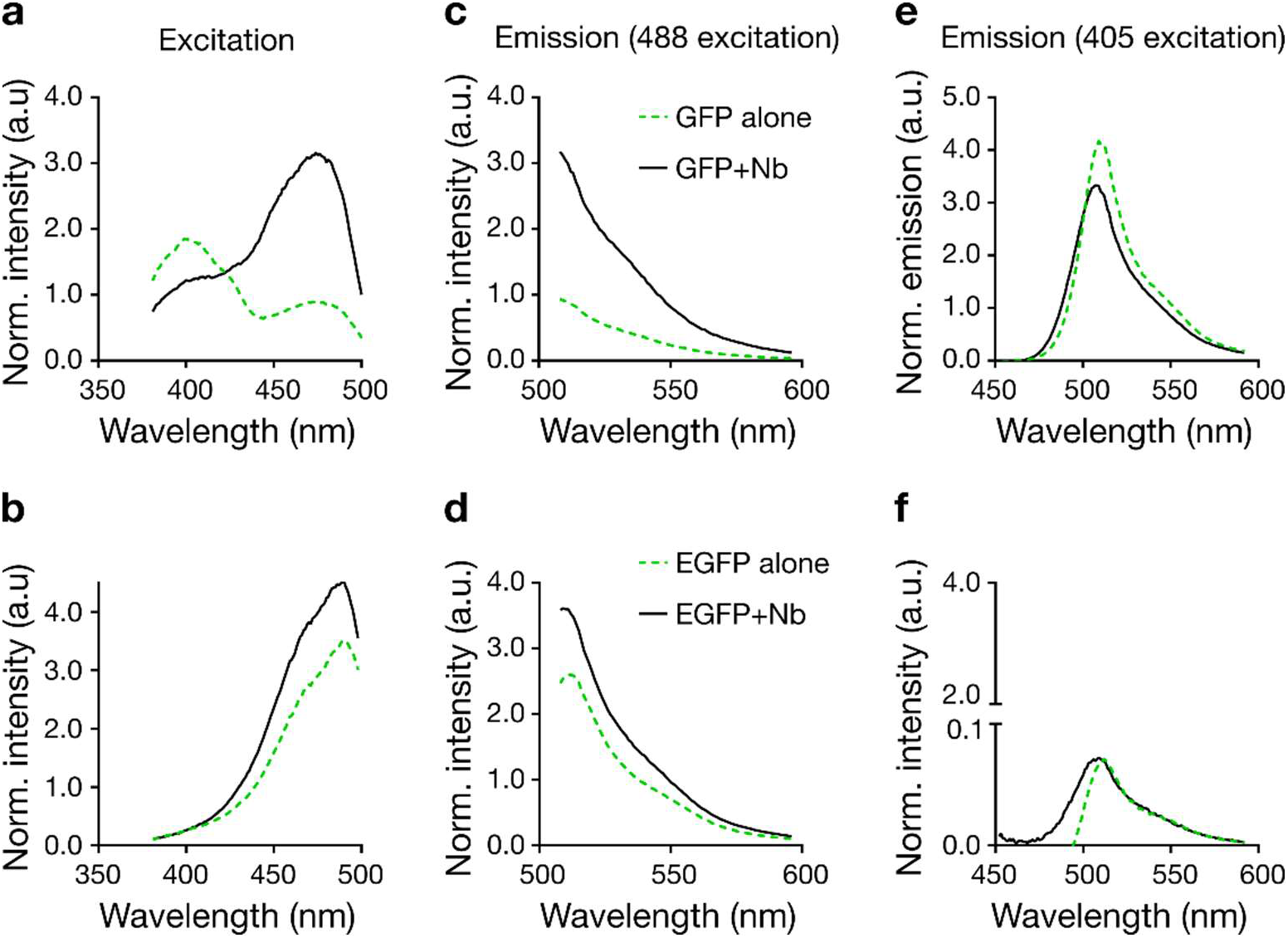
Change in excitation and emission spectra of recombinant GFP and EGFP in solution upon addition of unlabelled Nb. Excitation spectra for fluorescence detection at 510 - 520 nm (a,b) and emission spectra following 488 nm (c,d) and 405 nm excitation (e, f) of GFP-His (a,c,e) and EGFP-HIs (b,d, f) without Nb (green dashed line) and with unlabelled Nb (solid black line). All spectra are averages of three measurements.

### Nanobody binding on GUV membranes

In cell biology and microscopy, antibodies and nanobodies are commonly used to investigate the spatial organisation of membrane proteins. Therefore, we next tested changes in fluorescence properties of (E)GFP fluorescence at lipid membranes upon binding of unlabelled Nb. We first chose controlled conditions, employing GFP and EGFP attached to synthetic membranes of giant unilamellar vesicles (GUVs, made of dipalmitoylphosphatidylcholine (DOPC) lipid) via a His-tag (using DGS-NTA (1,2-dioleoyl-sn-glycero-3-[(N-(5-amino-1-carboxypentyl)iminodiacetic acid)succinyl])). For both GFP and EGFP, we observed an increase in total fluorescence intensity upon addition of unlabelled Nb, slightly higher (≈ 2 – 4 fold) than in solution and with slight differences between GFP and EGFP (4-fold compared to 2.5-fold, respectively) (Figure 2a,b).

**Figure 2.**
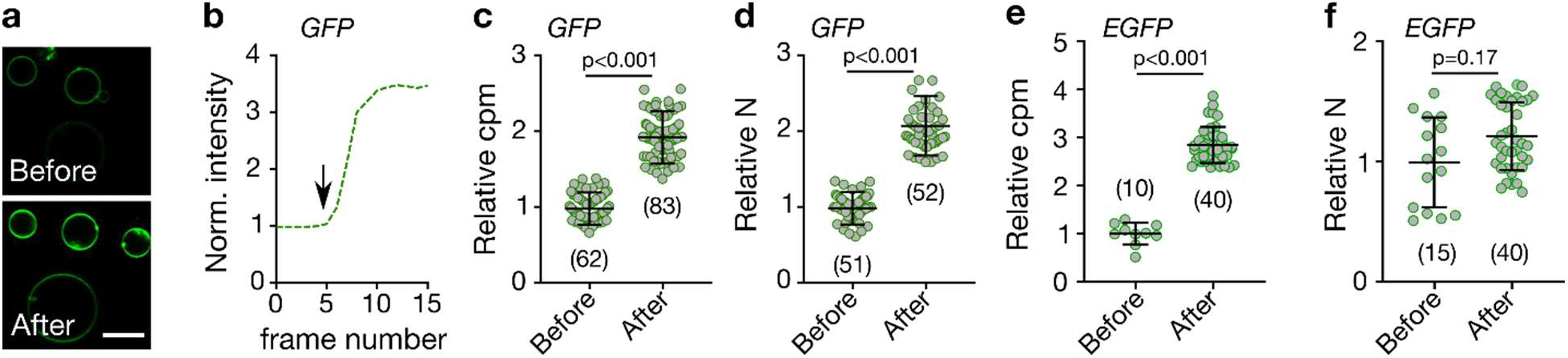
Effect of GFP-nanobody binding on GUV-anchored (E)GFP. Data for His-tagged (E)GFP anchored to GUVs (98 mol% DOPC and 2 mol% DGS-NTA) before and after addition of unlabelled Nb as marked. a) Representative confocal fluorescence microscopy images of the equatorial GUV plane for GFP. Scale bar 10 μm. b) Representative time-lapse over subsequently recorded confocal image frames of the normalized fluorescence intensity of the equatorial plane of a GFP-tagged GUV with time point of Nb addition marked with arrow. Relative change in (c) molecular fluorescence brightness (cpm) and (d) relative number of particles (N) of GFP before and after Nb addition as marked. Relative change in (e) cpm and (d) N of EGFP before and after Nb addition as marked. Values were determined from FCS experiments on individual (E)GFP-tagged GUVs. p-values were determined using the Kolmogorov–Smirnov non-parametric test. Number of data points is indicated on each graph.

To decipher this slight difference in increase in fluorescence intensity further, we tested how the fluorescence emission per individual GFP and EGFP molecule changed upon Nb binding. For this, we determined the molecular fluorescence brightness or fluorescence count rate per molecule, (cpm) derived from fluorescence correlation spectroscopy (FCS, Supplementary Figure 1a,b). FCS reveals the average emitted photons of a fluorophore by measuring photon statistics from multiple transits through the microscope’s observation spot. On GUVs, we observed an approximately 2-fold increase in molecular brightness of GFP following Nb addition (Figure 2c), which does not account for the ≈ 4-fold increase in total fluorescence signal intensity (Figure 2b). We therefore also derived the average number of fluorescing molecules, N, from the same FCS experiments (the amplitude of the correlation function. G(0) is inversely correlated to the average number of molecules N). Strikingly, upon Nb addition, we observed an approximately 2-fold increase in N (Figure 2d). For EGFP on GUVs there was also an approximately 2-fold change in molecular brightness cpm (Figure 2e) but in contrast to GFP only a marginal change in N (Figure 2f). As expected, for both GFP and EGFP the increase in N and cpm together account for the overall change in total fluorescence signal intensity (I = cpm × N).

While the increase in cpm upon Nb addition may be explained by the change in fluorescence excitation at around 480 nm as determined from the solution experiments (Figure 1a,b), the change in N and accordingly in concentration of fluorescing molecules suggests an unexplored enigmatic impact of Nb on the organisation of the membrane-bound GFP molecules. We will discuss this and the potential impact on assessing the spatial organisation of (E)GFP-tagged proteins in the plasma membrane of living cells in detail throughout the next sections.

### Nanobody binding on live-cell membranes

To test the effect of Nb on (E)GFP-tagged membrane protein organization further and in a more physiological setting, we next investigated the influence of Nbs on an GFP- and an EGFP-labelled glycosylphosphatidylinositol (GPI) anchored protein (GPI-AP) in the plasma membrane of living cells. Specifically, we expressed GFP-LYPD6 and GPI-EGFP in live PtK2 cells. GFP-LYPD6 is involved in Wnt signalling (Özhan et al., 2013), while GPI-EGFP is simply a lipid-anchored EGFP construct commonly used as probe to study GPI-AP organisation (Baumgart et al., 2016; Goswami et al., 2008; Saha et al., 2015; Schneider et al., 2017). We recorded confocal images (Figure 3a,b) as well as FCS data (Supplementary Figure 1c,d) to determine the total fluorescence intensity and values of cpm and N. For both proteins we found a modest increase in total fluorescence signal intensity upon Nb binding (≈1.1-fold for GFP and ≈1.5-fold for EGFP, Figure 3c,d), a slight increase in molecular fluorescence brightness (cpm, ≈1.1-fold for GFP and ≈1.5-fold for EGFP, Figure 3e,f), and a distinct variation in average number N of fluorescent molecules in the observation spot, with a slight ≈1.1-fold increase for GFP and a slight ≈1.2-fold decrease for EGFP (Figure 3c,f). However, especially the determination of the number of particles, N, was not straightforward on the live-cell membrane due to noise, cellular movements and spatial heterogeneity across the cells (i.e. due to variations in local concentrations as becomes obvious by variations in total intensity across one cell or between different cells, Figure 3 a,b). Overall, the impact of Nb binding on fluorescent protein tagged GPI-AP on living cells follows the trend from the model membranes with attenuated magnitude.

**Figure 3:**
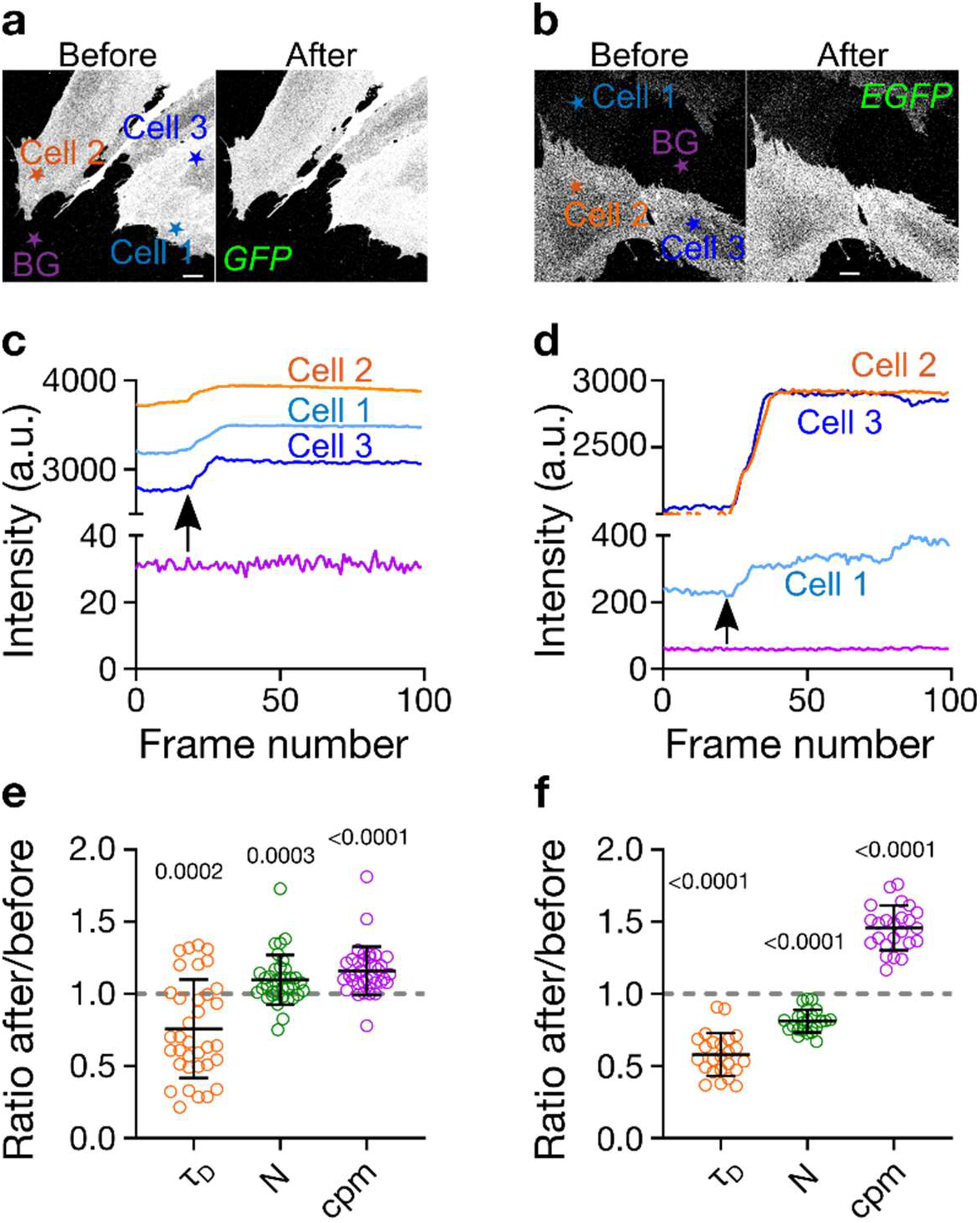
Effect of Nb-binding on (E)GFP in the plasma membrane of live PtK2 cells: GPI-anchored proteins GFP-LYPD6 (left panels) and GPI-EGFP (right panels). Representative confocal images before and after addition of Nb for a) GFP-LYPD6 and b) GPI-EGFP. Scale bars are 10 μm. Normalised fluorescence intensity traces for c) GFP-LYPD6 and d) GPI-EGFP for the cells as indicated in panel a and b, respectively (BG = background). Arrows show the time point when Nb was added. The intensities per frame represent mean values over each cell (see Materials and Methods for details). Change in τ_D_, N and cpm for e) GFP-LYPD6 and f) GPI-EGFP upon nanobody addition (values after Nb addition divided by values before). Change in average transit time (τ_D_ i.e. mobility), average fluorescing particle number (i.e. concentration, N), and molecular fluorescence brightness (cpm) upon Nb addition are determined from FCS experiments (one dot = one cell, for each cell 6 – 9 single FCS measurements were averaged, pooled data from three different days). The values on top of ratios in e,f indicate p-values obtained from Wilcoxon sign-rank non-parametric tests with hypothetical median values of 1 (ratio of 1 would indicate no change upon Nb addition).

### Nanobody effect on molecular mobility

So far, we obtained interesting insights from the stationary thermodynamic information from imaging and FCS (intensity, cpm and N), but the FCS measurements also allowed us to determine the average mobility of the membrane-anchored proteins. Measuring the diffusion dynamics can be performed robustly against local variations in concentration and expression levels and represents a way to study the organisation of plasma membrane constituents (Pinkwart et al., 2019; Schneider et al., 2020). From FCS, we obtained the transit time τ_D_, representing the average time it takes a molecule to cross the observation spot, and tested whether it changes upon Nb addition (Supplementary Figure 1). Intuitively, one may expect a slight decrease in mobility i.e. increase in values of τ_D_ upon addition of Nb due to the increased mass of the complex. Alternatively, one could expect no change at all as the mobility of membrane constituents is overwhelmingly determined by the properties of the membrane anchor (Saffman and Delbrück, 1975; Weiß et al., 2013). However, interestingly, we observed an increase in mobility after Nb addition for GFP-LYPD6 and GPI-EGFP in the membrane of live PtK2 cells (approximately 1.25 and 1.7-fold decrease in values of τ_D_ for GFP and EGFP, respectively; Figure 3e,f). From these τ_D_-values and the diameter d = 240 nm of the observation spot (full-width-at-half-maximum), we can estimate values of the diffusion coefficients given the diffusion equation 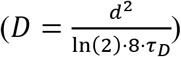 to D = 0.3 μm^2^/s and 0.4 μm^2^/s for GFP-LYPD6 without and with Nb and D = 0.8 μm^2^/s and 1.4 μm^2^/s for GPI-EGFP without and with Nb. Previously reported values for GPI-AP diffusion scatter from 0.3 to 1.0 μm^2^/s (Chojnacki et al., 2017; Eggeling et al., 2009; Huang et al., 2015; Lenne et al., 2006; Schneider et al., 2017; Veerapathiran and Wohland, 2017) where GPI-(E)GFP typically shows faster diffusion than other GPI-anchored probes (such as GPI-ACP or GPI-SNAP).

The apparent speed-up upon nanobody binding was puzzling, and to confirm these contradictory findings, we additionally recorded FCS data of GFP-LYPD6 and GPI-EGFP in living cells with higher statistical accuracy. Specifically, we performed scanning-FCS (sFCS) measurements, which yield simultaneous FCS data for multiple points along a quickly scanned line, i.e. hundreds of values of, for example, cpm and τ_D_ within a few measurement, which accounts for spatial heterogeneity and allows for the determination of average values with very high precision (Schneider et al., 2018; Waithe et al., 2018). The sFCS measurements confirmed the changes in values of τ_D_ i.e. faster diffusion for both GFP-LYPD6 and GPI-EGFP upon Nb binding (Figure 4a,b) in line with the point FCS measurements (Figure 3 e,f). Further, our sFCS data revealed that the increase in mobility (i.e. decrease in transit time τ_D_) was clearly correlated with an increase in brightness, cpm (Figure 4c-f), i.e. upon Nb addition the population of (E)GFP tagged GPI proteins shifted from a less bright and less mobile to a brighter and more mobile form.

**Figure 4.**
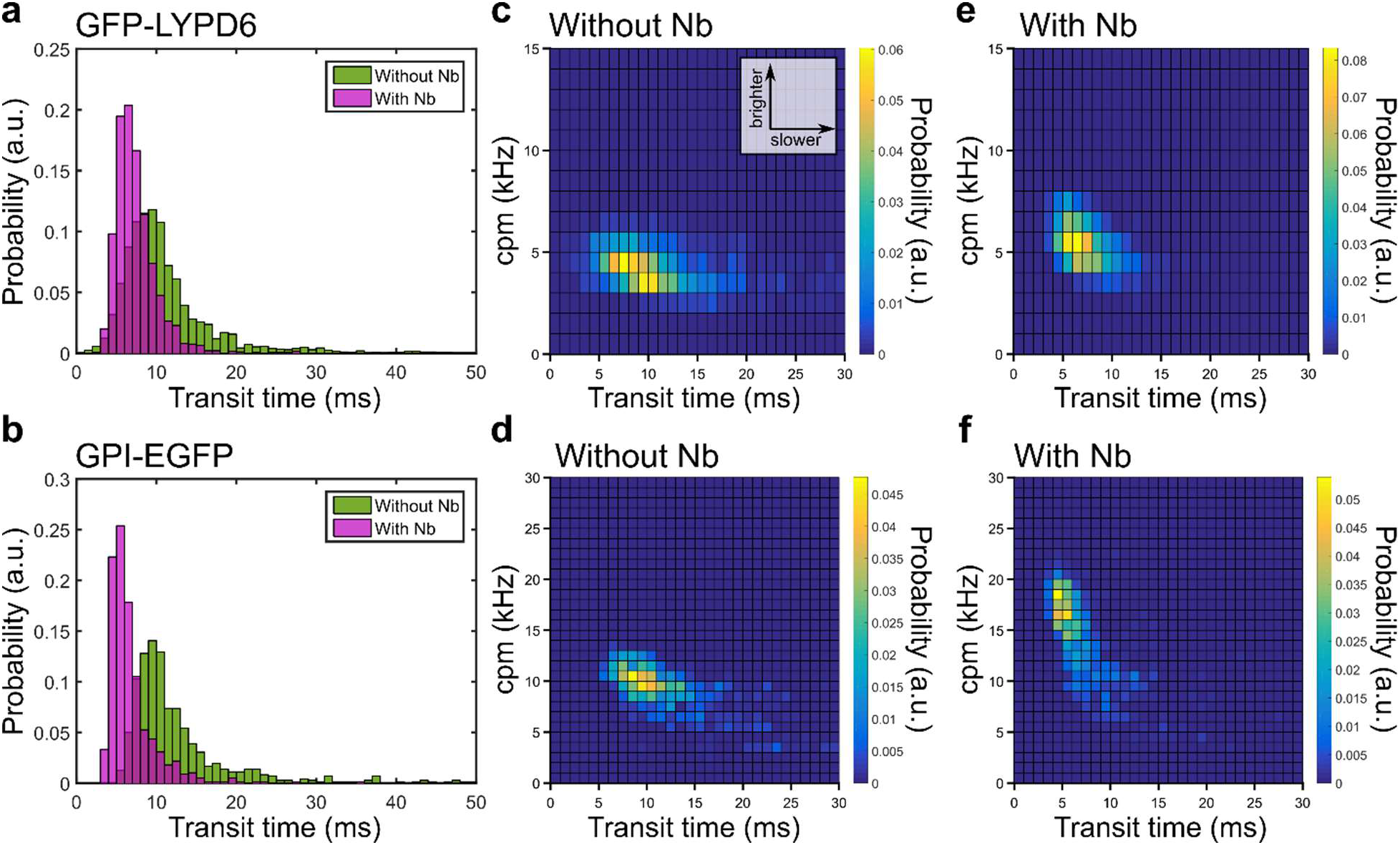
Effect of unlabelled Nb binding on mobility and brightness of GFP-LYPD6 and GPI-EGFP as probed by large sFCS datasets. Analysis of diffusion dynamics and molecular brightness (cpm) of PtK2 cells expressing GFP-LYPD6 (upper panels) and GPI-EGFP (bottom panels) and the effect of unlabelled nanobody. a,b) Transit time histograms for protein without (green) and with (magenta) presence of nanobody. >2000 curves single FCS curves for GFP-LYPD6 and > 700 for GPI-EGFP from >10 cells each. c-d) Two-dimensional pair value histograms (bi-variate histograms) of transit times and cpms for control (without Nb, c,d) and with addition of nanobody (e,f) for GFP-LYPD6 (top panels) and GPI-EGFP (bottom panels).

An explanation for these observations could be that the (E)GFP tagged proteins appear to a certain extend in aggregates or homo-oligomers that are (partially) disassembled after Nb binding, leading to an average increase in fluorescent particle number (Figures 2 and 3) and mobility (Figure 3 and 4) without majorly affecting the fluorescence lifetime (Supplementary Figure S2). GPI-EGFP dimers have been reported previously (Huang et al., 2015; Suzuki et al., 2012). However, since aggregates should in principle be brighter, one would in this case expect a decrease in molecular fluorescence brightness (cpm) upon aggregate disassembly. Our opposite observation indicates that the aggregates might actually be darker, e.g. due to self-quenching processes, and their fraction is rather low after disassembly. This is illustrated by the bi-variate histograms of cpm and τ_D_ (Figure 4c-f); the fraction of darker and slower molecules (“tail” of the distribution in Figure 4c,d) is notably reduced in the presence of nanobody (Figure 4e,f). If this is indeed the case, it is essential to know whether Nb binds to both oligo- and monomers with different affinity or selectively to one class.

### Diffusion of labelled nanobodies

To address what species is bound by the nanobody, we compared sFCS and FCS data of fluorescently labelled Nb and GFP-LYPD6 or GPI-EGFP at the plasma membrane of live PtK2 cells. Specifically, we employed Nbs tagged with the red-emitting dye Abberior Star 635P (AbStar635P-Nb), whose fluorescence emission was clearly distinguishable from that of the fluorescent proteins and detected on a separate detector. First, using confocal imaging we confirmed that the AbStar635-Nb bound only to the surface transfected cells and not to those without e.g. GPI-EGFP, i.e. AbStar635-Nb specifically interacted with the fluorescent protein on the membrane only (Supplementary Figure S3). Second, we also found an increase in mobility (i.e. decrease in average transit time τ_D_) and increase in brightness cpm of the EGFP tagged proteins upon AbStar635-Nb binding, i.e. the label did not influence this effect (Figure 5a,b and Supplementary Figure S4). Interestingly, simultaneously recorded sFCS data for AbStar635P-Nb and GPI-EGFP (Figure 5c,d) revealed a profoundly slower diffusion for AbStar635P-Nb compared to the EGFP-tagged proteins. We observed an average transit time of τ_D_ = 28.3 ms ± 1.0 (D = 0.5 ± 0.02 μm^2^/s) for AbStar635P-Nb and τ_D_ = 10.3 ms ± 1.0 (D = 1.2 ± 0.12 μm^2^/s) for GPI-EGFP (Figure 5a-d). This is an obvious contradiction, as the Nbs should be bound directly to the surface proteins (GPI-EGFP) but moved significantly slower than the protein itself. A possible explanation for this contradiction extends our previous hypothesis and points to the existence of two pools of (E)GFP on the cell surface; a darker oligomeric form that diffuses slowly and to which the Nb preferentially binds, which would drive the partial displacement of brighter and faster moving monomers, to which Nb does not bind efficiently.

**Figure 5.**
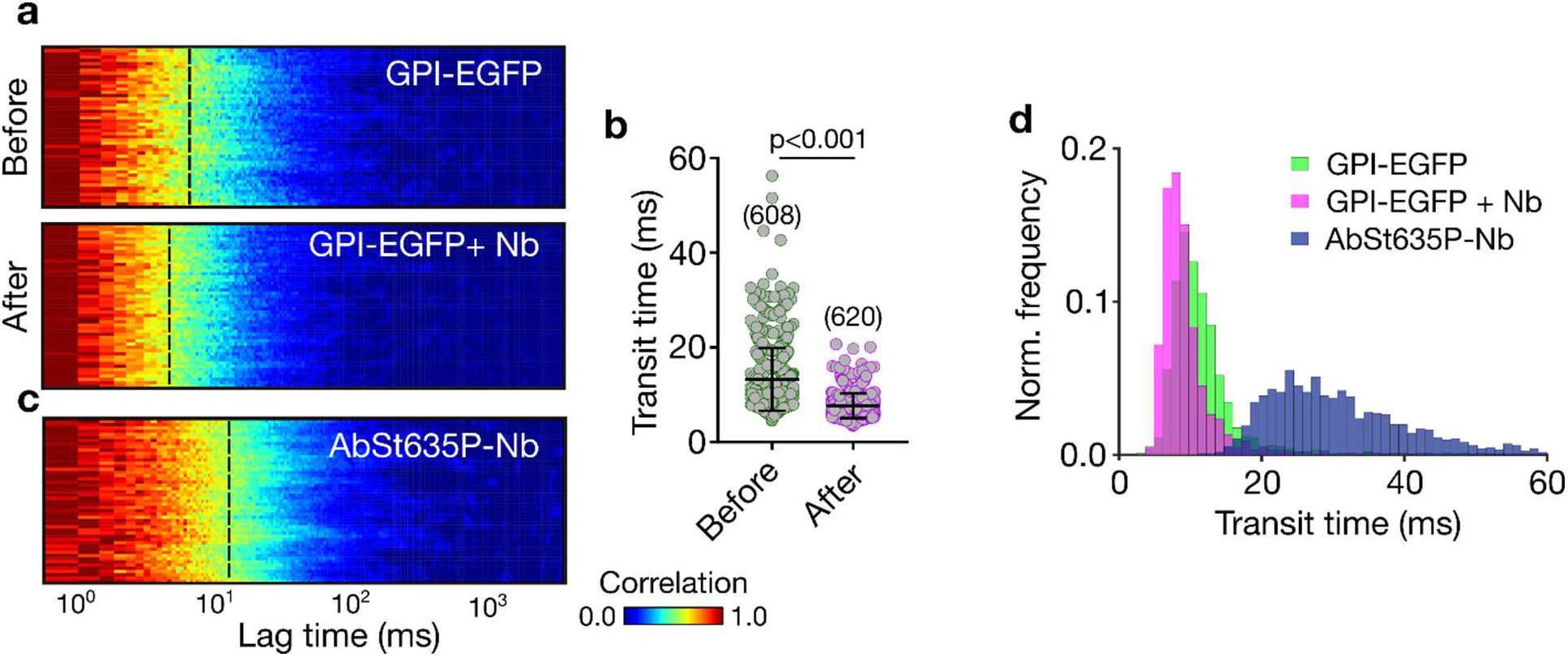
Diffusion of labelled Nb (AbStar635P-Nb) on GPI-EGFP expressing PtK2 cells. a,b) Representative normalised sFCS autocorrelation carpets for (a) GPI-EGFP before and after addition of AbStar635P-Nb (x: correlation lag time, y: line pixels (space), colour scale: normalised correlation from zero (blue) to one (red)), revealing a shift of the decay of the correlation curves (yellow region, average transit time highlighted by the dashed line) towards shorter times after addition of labelled Nb. b) τ_D_ values for GPI-EGFP before and after AbStar635P-Nb addition including mean values and standard deviations. c) Normalised autocorrelation carpet for AbStar635P-Nb bound to PTK2 cells expressing GPI-EGFP. d) Histogram of τ_D_ for GPI-EGFP (with and without Nb) and AbStar635P-Nb. The p-value given in panel b was calculated using the Kolmogorov–Smirnov non-parametric test.

### Missing co-diffusion of (E)GFP and nanobodies

To investigate these indications further, we applied fluorescence cross correlation spectroscopy (FCCS (Schwille et al., 1997)). Based on the principle of FCS, FCCS takes information from the temporal cross-correlation function of two simultaneously recorded fluorescence signal time traces of two distinctively labelled and emitting (e.g. green and red fluorescence, respectively) diffusing molecules to determine the degree of co-diffusion or interaction of the two molecules. Only when molecules show co-diffusion or interaction, the amplitude of the cross-correlation curve is larger than zero. An FCCS amplitude of zero indicates the absence of co-diffusion of the two fluorescing molecules (Bacia and Schwille, 2007). Therefore, the AbStar635P-Nb binding to fluorescently tagged GPI-APs should be a perfect sample for FCCS analysis, since every (red-emitting) Nb molecule should be bound to a (green-emitting) (E)GFP, yielding in theory a perfect cross-correlation between the red and green fluorescence signals. Performance of our FCCS experiments was validated through a positive control (red-labelled peptide binding specifically to membrane-embedded green-emitting cholesterol analogue, Supplementary Figure S5), showing a large non-zero FCCS amplitude, confirming near-perfect co-diffusion. Strikingly, we did not observe any notable cross-correlation and therefore no co-diffusion between AbStar635P-Nb and (E)GFP-tagged molecules anchored to the plasma membrane, as shown for GPI-EGFP in Figure 6a. The absence of co-diffusion is also illustrated by the dual-colour intensity time trace (Figure 6b) demonstrating only very rare detection events with signal from both channels, i.e. EGFP and AbStar635P-Nb independently crossed the observation spot. Following the same strategy as before, we used sFCS to confirm these findings for GFP-LYPD6, employing higher statistical throughput and spatial sampling. The auto-correlation carpets (Figure 6c,d) reveal the same slower diffusion of the bound AbStar635P-Nb compared to the binding partner GFP-LYPD6, as was the case for the experiments with GPI-EGFP (Figure 5 a,c). Similarly the scanning cross-correlation data of AbStar635P-Nb and GFP-LYPD6 confirmed the complete absence of co-diffusion of fluorescing binding partners and in contrast to a positive control (Supplementary Figure S6), we only observed noise (Figure 6e). We can only conclude that the Nbs do not bind the bright and fast diffusing (E)GFP-tagged proteins (supposedly monomers), but predominantly to a dark and slowly diffusing entity (supposedly oligomers).

**Figure 6.**
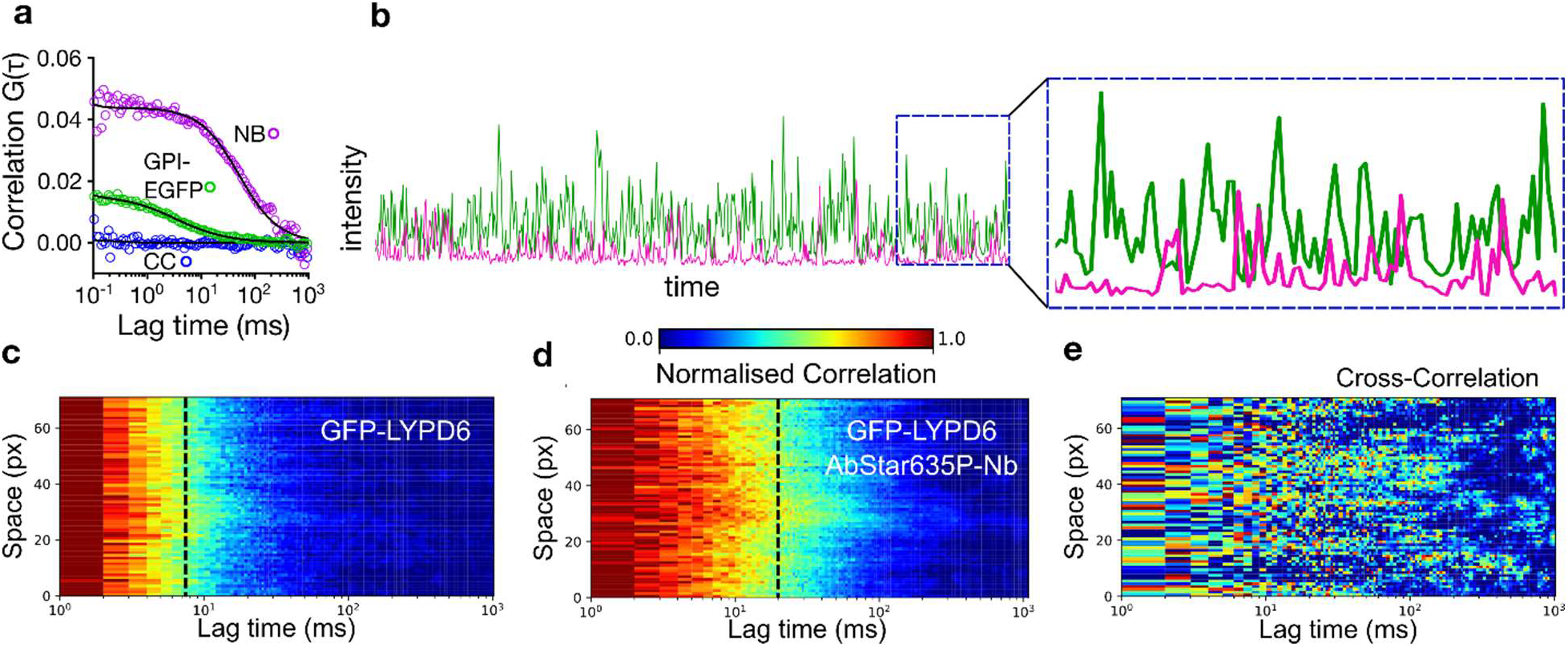
Missing co-diffusion of labelled Nb and (E)GFP-tagged surface proteins. PtK2 cells expressing GPI-EGFP or GFP-LYPD6 were treated with labelled Nb (AbStar635P-Nb) and FCCS data acquired. Positive cross correlation (CC) indicates interaction, i.e. co-diffusion. a) point FCCS of GPI-EGFP with autocorrelation EGFP (green), autocorrelation Nb (labelled with AbberiorStar635P, magenta) and CC (blue). b) Representative dual-colour intensity trace showing that the detection events for EGFP and AbStar635P-Nb rarely overlap in time. c-e) scanning FCCS data of GFP-LYPD6 expressed on the surface of PtK2 cells, with representative normalised auto-correlation data for (c) GFP-LYPD6, (d) AbStar635P-Nb and (e) normalised cross-correlation data of those. The dashed black line indicates the average transit times. The temporal cross-correlation of these two dataset does not show any positive cross-correlation, i.e. no co-diffusion.

Other reasons for the absence of cross-correlation could be i) very fast binding kinetics (i.e. on- and off-rates) of the Nb-(E)GFP interaction or ii) too low binding leaving too many unbound (E)GFP molecules. However, our data do not support these scenarios. 1) We determined off-rates, k_off_, for the Nb-GFP binding using surface plasmon resonance (SPR) experiments (of GFP binding to surface-immobilised (labelled and unlabelled) Nbs), resulting in k_off_ ≈ 5 × 10^-4^ s^-1^ (Supplementary Figure S7). Consequently, the Nb-(E)GFP complex is stable for about 30 minutes, which is in agreement with previous data (Della Pia and Martinez, 2015). During the 1-50 milliseconds long transit through the observation spot, the Nb-(E)GFP complex should be intact. This data rules out fast kinetics. 2) We also recorded FCCS data in large excess of AbStar635-Nb, saturating GFP binding. However, we still did not observe any cross-correlation (Supplementary Figure S8). This data rules out the domination of unbound particles.

We also tested whether the lack of cross-correlation signal could be dye-specific for AbStar635P, but cross-correlation was also absent when performing the same sFCCS experiments as before with nanobody labelled with the dye Atto594 (Supplementary Figure S9).

Another possible explanation for the lack of co-diffusion might be an extremely efficient (close to 100%) energy transfer (Sun et al., 2011) between the fluorescent protein(s) and the AbStar635-Nb. Such a Förster resonance energy transfer (FRET) would render the Nb-bound fluorescent proteins very dim and therefore hardly visible for FCCS analysis (FCS-based experiments require rather large fluorescence brightness (Saffarian and Elson, 2003; Schneider et al., 2020)). Several observations oppose this scenario: i) Close to 100% FRET efficiency should lead to a huge decrease in the number N of visible donor (E)GFP molecules and thus a decrease of EGFP fluorescence intensity with nanobody binding, which is not the case (Figure 1, 2 and Supplementary Figure S10). ii) A large energy transfer generally leads to a vast decrease in the fluorescence lifetime of the FRET donor, in this case (E)GFP. We therefore measured and compared values of the fluorescence lifetime for GFP and EGFP with and without binding to unlabelled and AbStar635P-labelled Nb. There was a small reduction in fluorescence lifetime for GFP (≈ 3 ns to ≈ 2.2 ns) and EFGP (≈ 2.6 ns to ≈ 2.1 ns) in solution (Supplementary Figure S11), indicating only a minor influence by FRET and not explaining the complete lack of cross-correlating fluorescence signal. Some FRET may explain though the lower increase in fluorescence intensity upon labelled nanobody binding to the recombinant proteins in solution (Supplementary Figure 12) compared to unlabelled Nb (Figure 1). iii) The molecular fluorescence brightness cpm of the donor EGFP molecules should go down after addition of labelled compared to unlabelled Nbs. As highlighted before, we however see an increase in cpm values independent of labelled or unlabelled Nb (Figure 4c-f and Supplementary Figure S4).

## Conclusions

The use of Nbs, not only as an alternative for full-length antibodies, presents a versatile new route for detection and manipulation of proteins in biology and especially microscopy (Beghein and Gettemans, 2017). Their small size, monovalancy, the ability to label them stoichiometrically, and their recombinant or even in vivo production makes them an attractive tool (Bothma et al., 2018; Grußmayer et al., 2014; Sograte-Idrissi et al., 2020). The GFP-binding Nbs along with Nbs against other fluorescent proteins enable conveniently to perform super-resolution microscopy on the tagged protein of interest (Platonova et al., 2015; Ries et al., 2012; Sograte-Idrissi et al., 2019). A modulation of the GFP’s spectral properties by interaction with Nbs has been shown and exploited before (Bothma et al., 2018; Kirchhofer et al., 2010), however the Nb has not been implicated in the reorganisation of the tagged protein.

In this study, we used imaging and spectroscopic techniques to investigate changes in the dynamic organisation of (E)GFP tagged lipid anchors in model membranes and living cells upon binding of labelled and unlabelled anti-GFP nanobody. Overall, our data show that (i) Nb increases the apparent number of GFP molecules on GUVs and cells, (ii) Nb binding to fluorescent proteins on living cells increases the mobility of GPI-anchored EGFP or GFP, (iii) the Nb diffuses slowly compared to the GFP on living cells, which means that they do not co-diffuse, and (iv) the Nb binding modulates photophysical properties of the fluorescent proteins. These findings may suggest that the Nbs bind predominantly to an already existing dark form of the (E)GFP, which might be a slow-moving, higher order complex. Lack of co-diffusion suggests that this complex is not strongly fluorescent (e.g., due to self-quenching) but still primarily recognized by the Nb. Upon binding, Nb could partially disassemble the dark complex and release a few molecules from the complex that become fluorescent but are not bound to a Nb. Although this process still needs to be directly shown in the future, our results imply that the Nb binding could influence the organization of the GFP-tagged proteins in living cells. It has been reported before that Nb binding could perturb protein function (Küey et al., 2019). Consequently, when performing conventional or super-resolved imaging using anti-GFP nanobodies to label (E)GFP conjugated proteins, the measurements need to be interpreted with care, especially on live cells and when quantitative data on dynamics are derived.

We only showed the effect of GFP-nanobodies, thus we refrain ourselves from generalizing the effect to all nanobodies. Recently, by using SNAP-25 and Syntaxin 1A nanobodies, a previously undetected pool of synaptic population were found in the cells (Maidorn et al., 2019), which was attributed to nanobodies’ ability to reveal different organization patterns. Therefore, there is a possibility that the effect of nanobodies could be more general. Moreover, here we only test nanobodies against GFP proteins (in model membranes) or GFP-labelled GPI-anchored proteins, but it seems probable that the effect will be similar on other proteins labelled with GFP or its derivatives as the modulating interactions are between the fluorescent protein and the nanobody.

## Significance

Nanobodies, especially against fluorescent proteins, are widely used in microscopy to investigate the organisation of recombinantly tagged proteins. Usually, the fluorescent label on the nanobody is imaged as proxy for the organisation of the protein of interest. Here, by applying imaging and single molecule fluorescence spectroscopy, we show that in live cells, the distribution and dynamics of nanobody and target protein of interest may differ. We used GPI-anchored proteins as an example and illustrate that the nanobody bound to GPI-EGFP did not accurately resemble the native organisation of the protein and may even change it. Since nanobodies are constantly becoming more popular, our findings are crucial as they suggest that it is necessary to exercise prudence when applying nanobodies for quantitative analysis of live-cell microscopy.

## Acknowledgements

We thank the EMBL Super-resolution course, Dr. Marco Lampe and Dr. Jonas Ries for their help on emergence of this interesting question, Dr. Chris Paluch for his help on Biacore SPR analysis. We also thank Christoffer Lagerholm and the Wolfson Imaging Centre Oxford and the Micron Advanced Bioimaging Unit (Wellcome Trust Strategic Award 091911) for providing microscope facility and financial support. We acknowledge funding by the Wolfson Foundation, the Medical Research Council (MRC, grant number MC_UU_12010/unit programmes G0902418 and MC_UU_12025), MRC/BBSRC/EPSRC (grant number MR/K01577X/1), the Wellcome Trust (grant ref 104924/14/Z/14), the Deutsche Forschungsgemeinschaft (Research unit 1905 “Structure and function of the peroxisomal translocon”, Jena Excellence Cluster "Balance of the Microverse", Collaborative Research Center 1278 "Polytarget"), Oxford-internal funds (John Fell Fund and EPA Cephalosporin Fund) and Wellcome Institutional Strategic Support Fund (ISSF). ES is funded by the Newton-Katip Celebi Institutional Links grant (352333122) and SciLifeLab fellowship.

## Author contributions

F.S. and E.S. designed the study, performed the experiments and analysed the data. F.S., E.S. and C.E. wrote the manuscript.

## Materials and Methods

### Cell culture & labelling

PtK2 cells were cultured at 37 °C, 5 % CO_2_, in DMEM (Sigma Aldrich) supplemented with 16% FBS (Sigma Aldrich). For microscopy the cells were seeded on 25 mm diameter glass coverslips (#1.5 thickness). Transfections of GPI-EGFP (kind gift from Kai Simon’s lab) and GFP-LYPD6 (Özhan et al., 2013) were performed with Lipofectamine 3000 (Thermo Fisher) according to the manufacturer’s protocol.

While imaging cells at 37 °C, they were incubated with unlabelled (GFP-binding protein, Chromotek) and Abberior 635P conjugated Nanobody (GFP-booster, Chromotek). All experiments were performed in L15 imaging medium (Sigma Aldrich) and the nanobody added to 1 mL and well mixed.

### GUVs

GUVs were prepared using electro-formation as described in(Jenkins et al., 2019). Lipid mixture (1mg/mL DOPC:DGS-Ni-NTA (both obtained from Avanti Polar Lipids) 96:4 molar ratio) were spread on platinum wire and dipped into 300 mM sucrose. GUVs formed after exposure to an AC filed of 2 V and 10 Hz for 1 h followed by 2 V 2 Hz for another 30 minutes. His-tagged GFP (Sino Biological) and eGFP (OriGene) was incubated for 20 minutes with the vesicles before imaging and FCS was performed.

### Confocal microscopy & FCS

Confocal microscopy and FCS were performed on a Zeiss780 and Zeiss880 both equipped with an Argon laser for fluorescence excitation. All sFCS and most imaging has been performed in photon counting mode using Channel S. To excite the labelled nanobody the HeNe 633 excitation has been employed. For single colour FCS and imaging a 488 dichroic mirror and for two-colour imaging a 488/561/633 MBS was used. The fluorescence was collected between 500 nm and 600 nm for the green channel and between 640 nm and 695 nm for the red channel. Laser powers were between 1 and 5 μW and kept below saturation to avoid artefacts in FCS.

The images were processed using FIJI (Rueden et al., 2017; Schindelin et al., 2012). The plasma membranes of each cell was segmented out using the polygon selection tool. Similar sized regions of interest were generated for the background. The average intensities over time for the whole area were extracted from the z-profile.

Point-FCS measurements were performed using Zeiss’ internal FCS routine. Measurements were between 10 and 15 seconds long. The objective’s correction collar was adjusted and the pinhole aligned measuring the diffusion of Alexa Fluor 488 in water. FCS measurements were saved as .fcs files for fitting. The same procedure was followed for cross-correlation (FCCS) measurements but additionally using a cross-correlation positive control (Bodipy and Alexa647 labelled HDL particles (Plochberger et al., 2017)) to ensure optical alignment.

sFCS measurements were performed as xt scans. 52 pixels were acquired for 10^5^ lines at about 2000 Hz yielding pixel dwell time of 3.94 μs (overall resulting in a total acquisition time of about 47 seconds). The data were saved as .lsm5 file and externally correlated using the FoCuS_scan software package (Waithe et al., 2018). To correct for photobleaching, the first seconds were cropped off and a local averaging bleaching-correction applied as described in (Waithe et al., 2018). sFCCS measurements were performed in a similar manner using the described acquisition settings in conjunction with the optical set-up for two-colour imaging as described above. As a positive control for sFCCS, we used a sparse sample of DOPC (1,2-dioleoyl-sn-glycero-3-phosphocholine), Avanti Polar Lipids) vesicles doped 1:50,000 with DiO (Invitrogen) and 1:10,000 with AbberiorSTAR-Red-PEG-Cholesterol (Abberior). The sample was prepared by mixing the lipid and dyes in ethanol, drying the mixture, and re-suspending the lipid film by vortexing and ultra-sonication (5 minutes and 30 minutes, respectively) in water. Measurements were performed in PBS.

Point and sFCS data were fitted in FoCuS (Waithe et al., 2016, 2018). Point-FCS data showed a contribution of a triplet component (40 μs for (E)GFP (Schneider et al., 2020)), a fast (probably cytoplasmic) component, with transit times around 0.1 ms, and a slower transit time which was attributed to the diffusion in the membrane. Thus, pFCS data were fitted with a two component diffusion model including a triplet state (Widengren et al., 1995). sFCS acquisitions miss the fast dynamics and were fitted with a single component 2D diffusion model (Schneider et al., 2018). Fitting in FoCuS is performed using a Levenberg-Marquard non-linear least suare optimisation. The fitted parameters including the cpms (determined from fitted amplitude and the average count rate) were exported as Excel sheets and post processed with Matlab (Schneider et al., 2018).

Some data on mobility were acquired cell by cell to account for the inherent biological heterogeneity. In these cases only the ratio of the transit times, number of molecules or counts per molecule before and after addition of the nanobody are reported (After/Before). Statistical tests were performed in GraphPad Prism 8. We employed the Wilcoxon sign-rank non-parametric tests with hypothetical median values of 1 for the data presented as ratios and we used the Kolmogorov– Smirnov non-parametric test for all other data.

### Lifetime measurements

Life time imaging was performed on a Microtime 200 (PicoQuant) equipped with a FlimBee galvo scanner. Fluorescence was excited with a 488 nm diode laser (PicoQuant) and focused onto the sample with an Olympus UPlanSApo 60 x water-immersion objective. Images were acquired as 50 by 50 μm^2^ (512 by 512 pixel) for collecting the fluorescence for 60 s at low excitation power (<1 μW) to avoid too high count rates and distortion of the TCSPC data (Isbaner et al., 2016). The overall TCSPC curves were used for lifetime fitting (2 component tail fit, second component fixed to 4.1 ns). We report the amplitude weighted lifetime of GPI-eGFP and GFP-LYPD6 in the plasma membrane of living PtK2 cells.

The same set-up was used to measure the fluorescent lifetime of GFP-His and EGFP-His in solution. The data were acquired as point scans.

### SPR

We immobilised the nanobodies (either fluorescently labelled or unlabelled) by amine coupling to a CM5 chip (with an RFPNB in the reference channel) then injected GFP as the analyte in a kinetic analysis (a single injection at 87 nM, using curve fitting in the BiaEvaluation software to measure on and off rates from which the K_D_ is calculated).

### Spectra

All spectral measurements were performed using a CLARIO STAR plate reader (BMG LABTECH). His-tagged GFP and EGFP were measured in PBS in glass bottom 96-well plates (Porvair Sciences) which were prior to the measurements coated with BSA to prevent sticking of the fluorescent proteins to the glass. All spectra are averages of multiple wells. Excitation scan were performed with a readout at 510 nm and emission scans were performed with excitation at 405 nm or 488 nm. We choose a spectral resolution of 1 nm and used 200 flashes per wavelength for averaging.

