## Supplementary Figures for "Influence of nanobody binding on fluorescence emission, mobility and organization of GFP-tagged proteins"

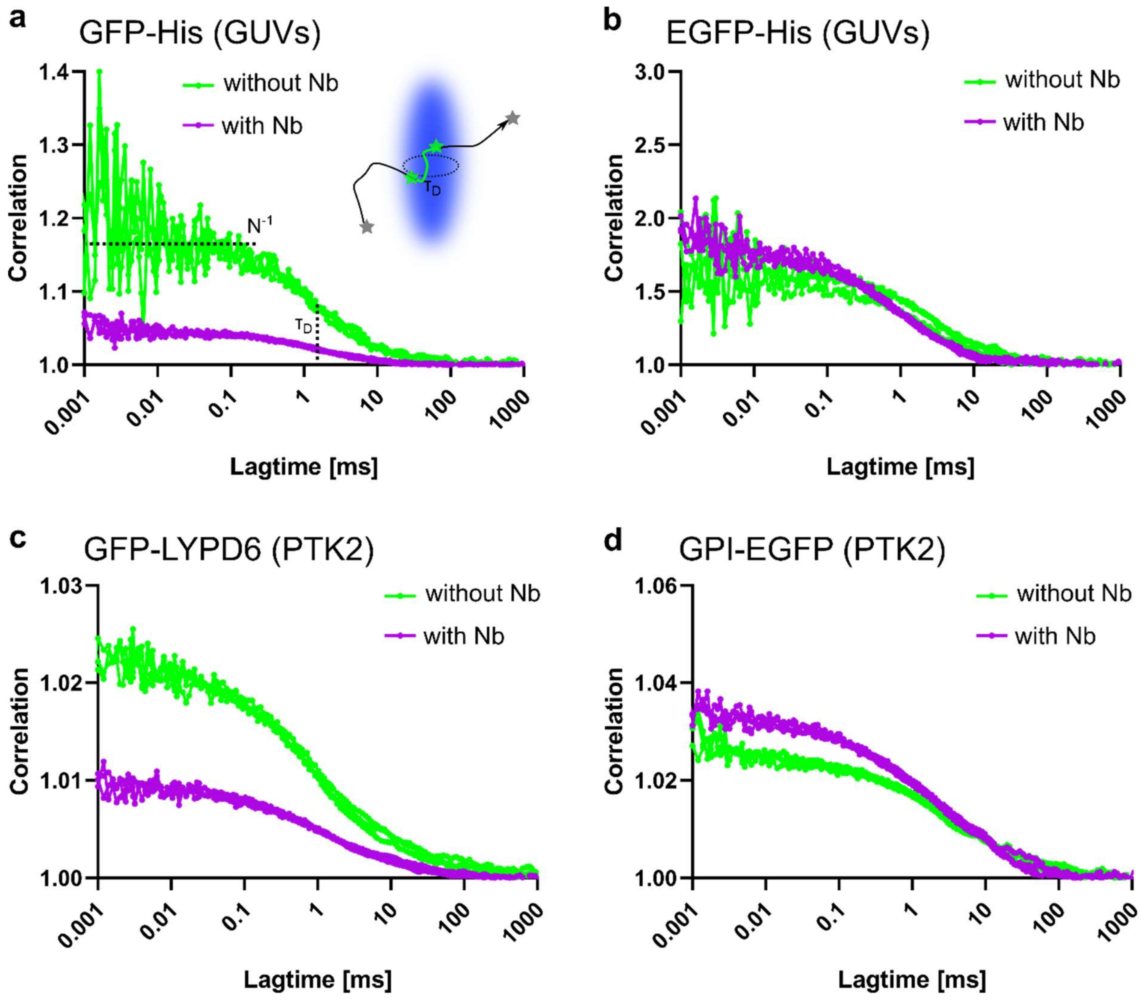


**Supplementary Figure S1: Representative point FCS data**. GFP (left panels) and EGFP (right panels) diffusing on different membrane systems, GUVs (98 mol% DOPC and 2 mol% DGS-Ni-NTA) decorated with GFP-His (a) and EGFP-His (b), and plasma membrane of live PtK2 cells for GFP-LYPD6 (c) and GPI-EGFP (d) with (magenta) and without (green) addition of unlabelled Nb; three representative curves are shown for each condition. Inset of a) schematic of FCS measurements for the determination of the average molecular brightness or count-rate per molecule (cpm), the average number of fluorescing molecules in the observation spot (N), and the average transit time $\tau_D$ through the observation spot (blue) as measure of the molecular mobility.

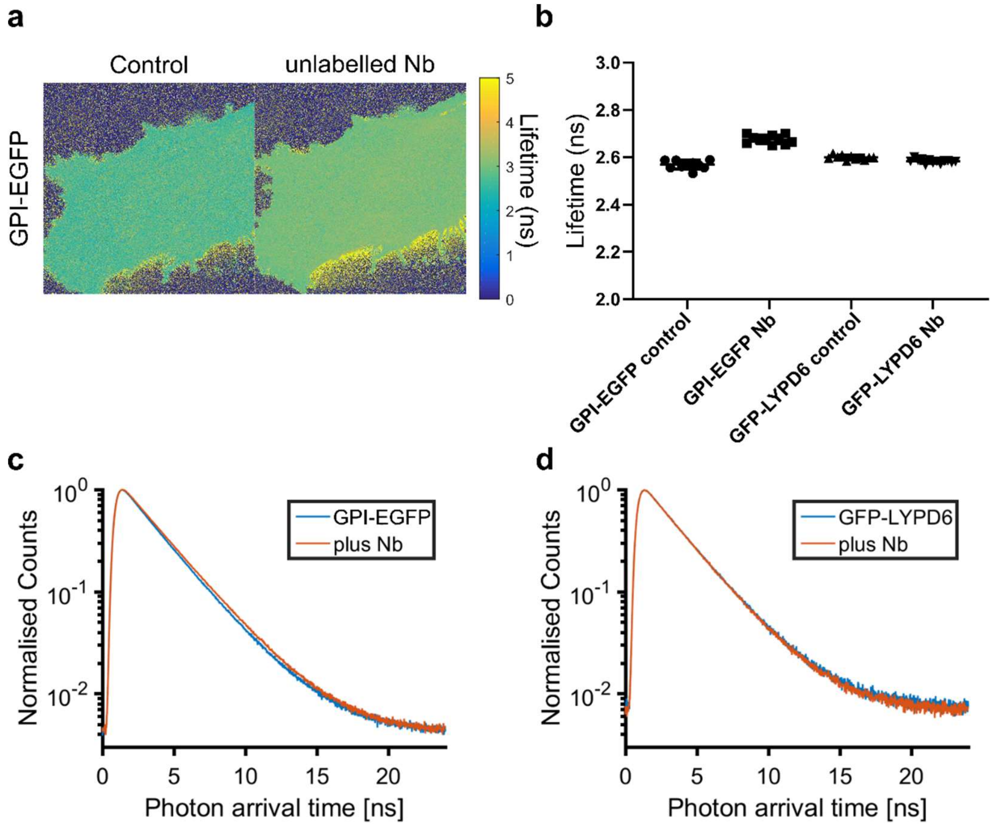


**Supplementary Figure S2: Fluorescence lifetime imaging of GPI-EGFP and GFP-LYPD6.** a) Representative lifetime images of the basal plasma membrane of live PtK2 cells transfected with GPI-EGFP (left) and treated with unlabelled Nb (right). Image size 50x50 $\mu m^2$. b) Average values of fluorescence lifetimes of the same fluorescent proteins as determined from fitting the respective TCSPC-based fluorescence decays averaged over the whole image (c,d). Note that amplitude weighted lifetimes from a bi-exponential tail fit are given as lifetimes for both GPI-EGFP and GFP-LYPD6.

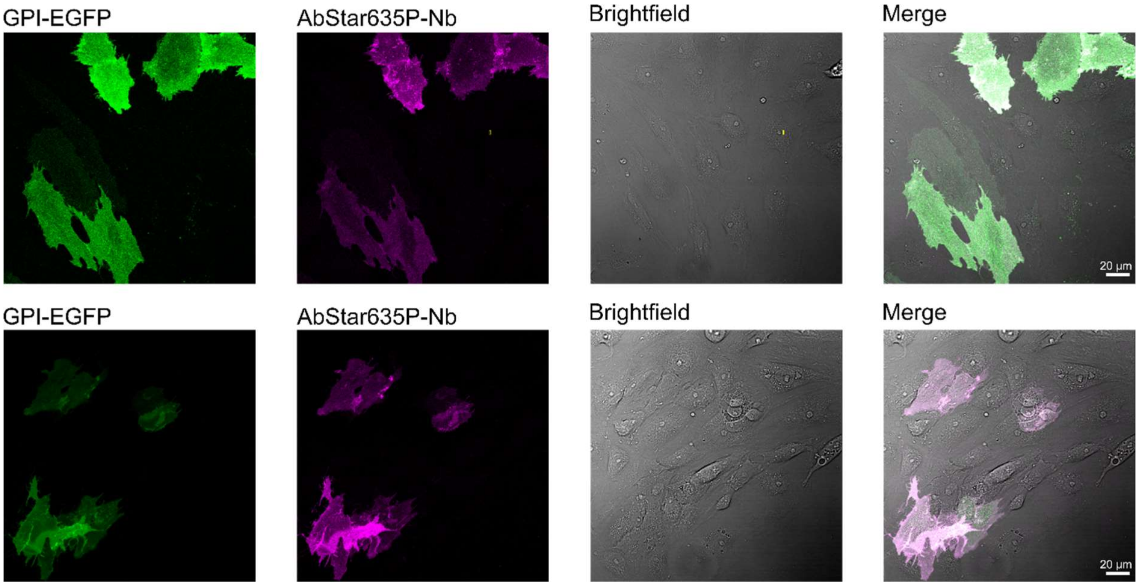


**Supplementary Figure S3: Specific binding of the labelled nanobody (AbStar635-Nb) to transfected cells.** Representative confocal images of PtK2 cells expressing GPI-EGFP and additionally stained with Abberior Star 635P-labelled nanobodies (AbStar635P-Nb): EGFP (left), AbStar635P-Nb (middle left), brightfield transmission (middle right) and merged (right) observation channels. Scale bars 20 µm.

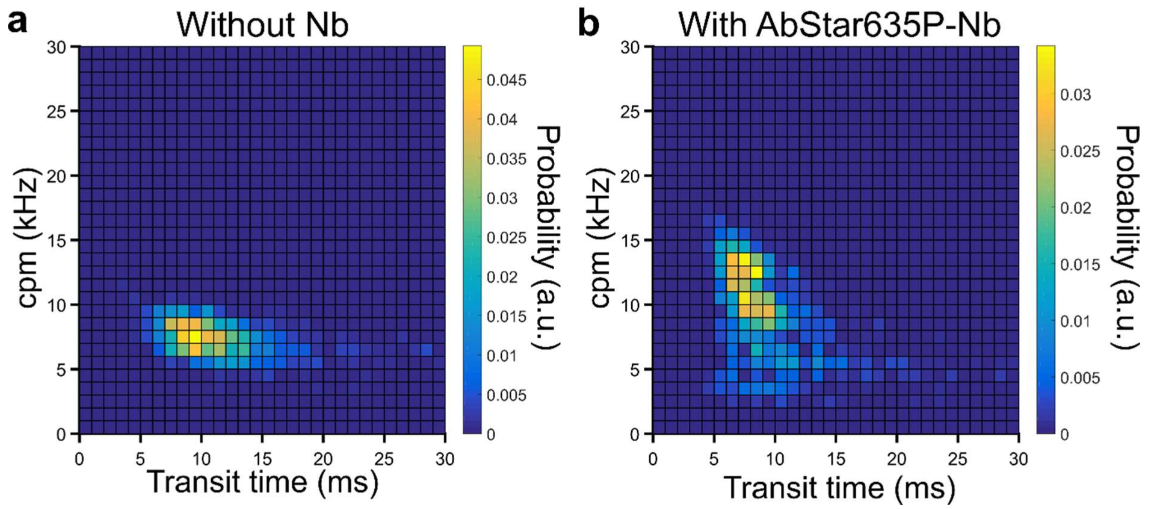


**Supplementary Figure S4: Also labelled Nb causes shifts in mobility and brightness.** Two-dimensional pair value histograms (bi-variate histograms) of transit times and cpms from sFCS experiments on GPI-EGFP on live PtK2 cells without (a) and with a labelled Nb (AbStar635P, b). The observed shifts are similar to the unlabelled Nb in Figure 4d,f.

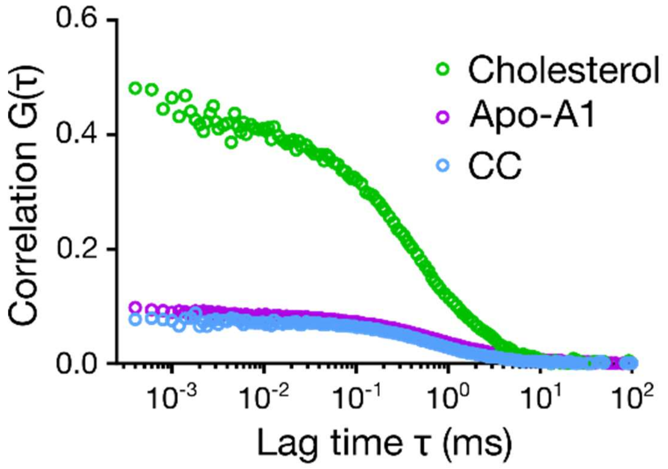


**Supplementary Figure S5: FCCS positive control.** Representative FCCS data (autocorrelation (green and magenta) and cross-correlation (CC, blue) curves) for HDL particles labelled with Bodipy-cholesterol and ApoA1-Alexa647, demonstrating the capability of the set-up to obtain almost 100% cross-correlation for a perfect sample.

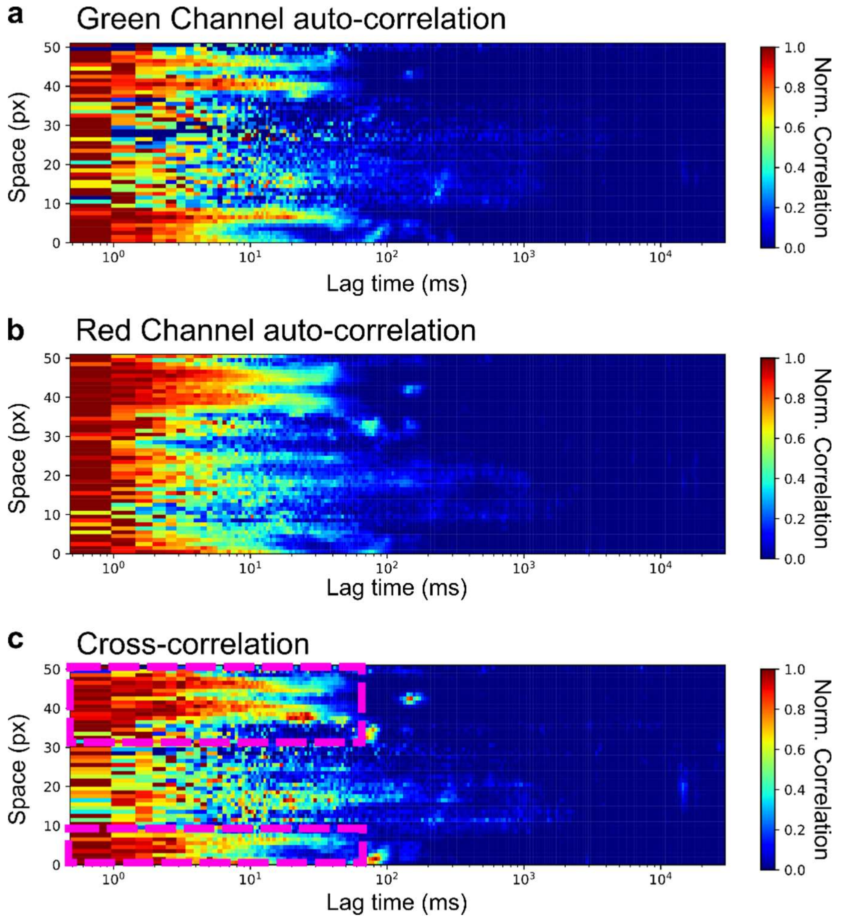


**Supplementary Figure S6: sFCCS positive control.** Exemplary sFCCS measurement of a sparse vesicle solution. Vesicles were made of DOPC doped with DiO and AbberiorSTAR-Red-PEG-Cholesterol. Normalised auto-correlation for the green (a, Fast-DiO) and red channel (b, AbberiorSTAR-Red), and cross-correlation of both (c). The dashed magenta boxes in c) indicate positions of clear cross-correlation resembling the respective auto-correlations and revealing co-diffusion.

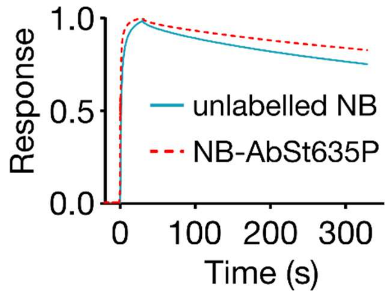


**Supplementary Figure S7: Nanobody and GFP interaction kinetics.** Surface plasmon resonance (SPR) plots showing the strong (and fast) binding of His-GFP to immobilized unlabeled (blue) or Abberior Star 635P tagged Nb (dashed orange) suggesting that the complex should be stable over the course of the experiments ($k_{off}$= 5.554 $e^{-4}$ $s^{-1}$ and $k_d$=3.8 $e^{-11}$ M).

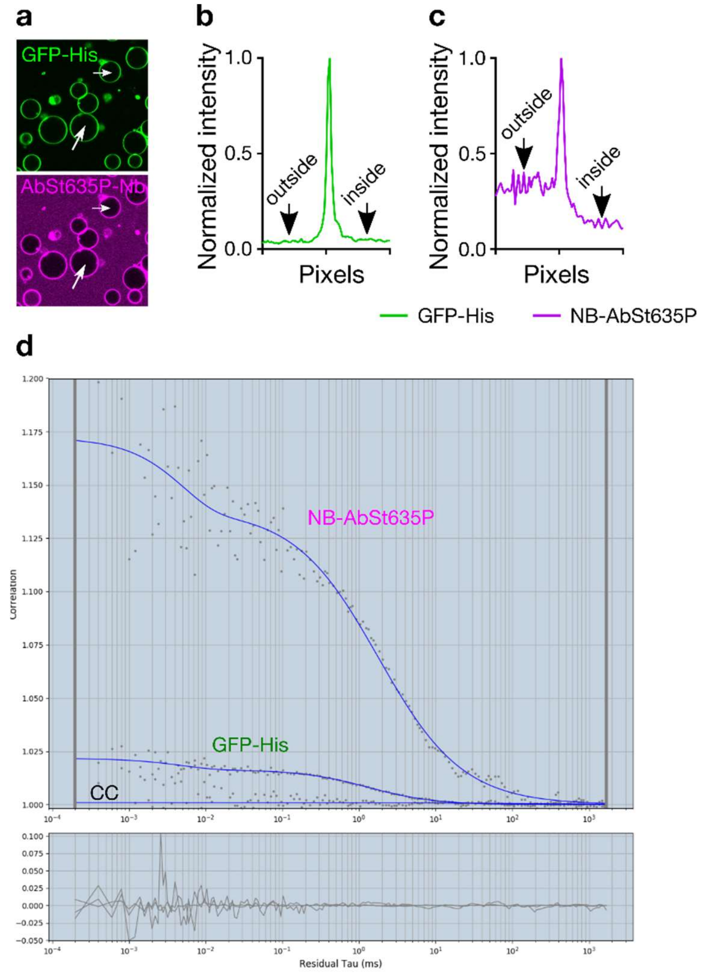


**Supplementary Figure S8: FCCS measurements at excess Nb**. a) Confocal images of GUVs decorated with His-GFP and incubated with AbStar635P-Nb (80 µm x 80 µm). b, c) The line profile of the arrows shown in panel a for b) GFP and c) AbStar635P-Nb. The image and the line profile show excess Nb in the solution. d) Autocorrelation and cross-correlation for GFP and AbStar635P-Nb. No cross-correlation is observed when performing point FCCS measurements on the GUVs.

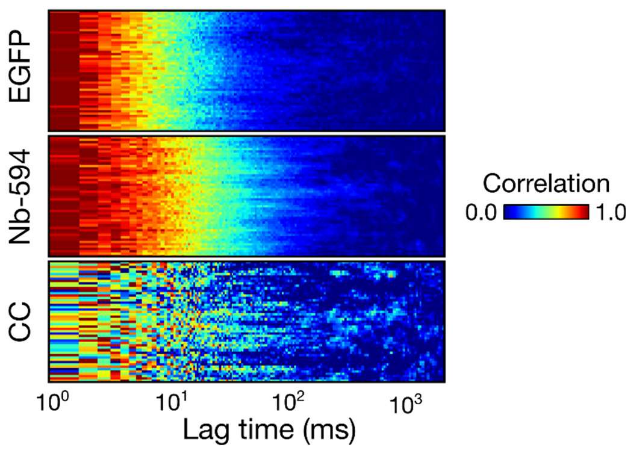


**Supplementary Figure S9: No influence of the label on the nanobody.** Representative cross-correlation is also absent using a nanobody labelled with Atto594. sFCCS measurement on GPI-EGFP transfected PtK2 cells additionally tagged with a labelled nanobody (labelled with Atto594). The nanobody channel resembles the familiar slow-down but only neglectable cross-correlation (CC) can be observed.

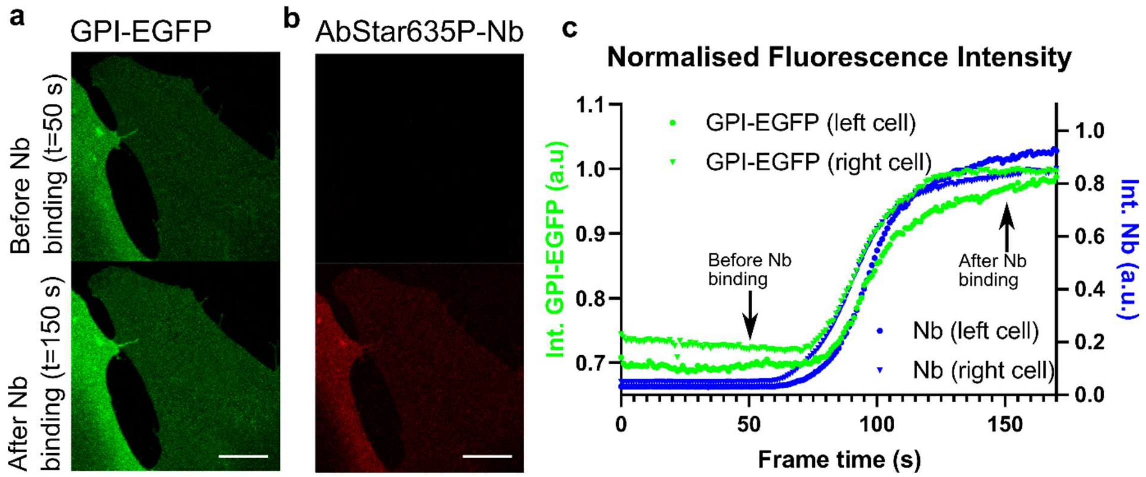


**Supplementary Figure S10: Changes in fluorescence intensity upon addition of AbStar635P-Nb to live PtK2 cells.** a,b) Representative confocal images of the basal plasma membrane of a PtK2 cell expressing GPI-EGFP (green) before and after addition of AbStar635P labelled Nb (red) taken from an image stack at t=50s and t=150s after addition. c) Extracted and normalized fluorescence intensity time traces for both cells in a,b.

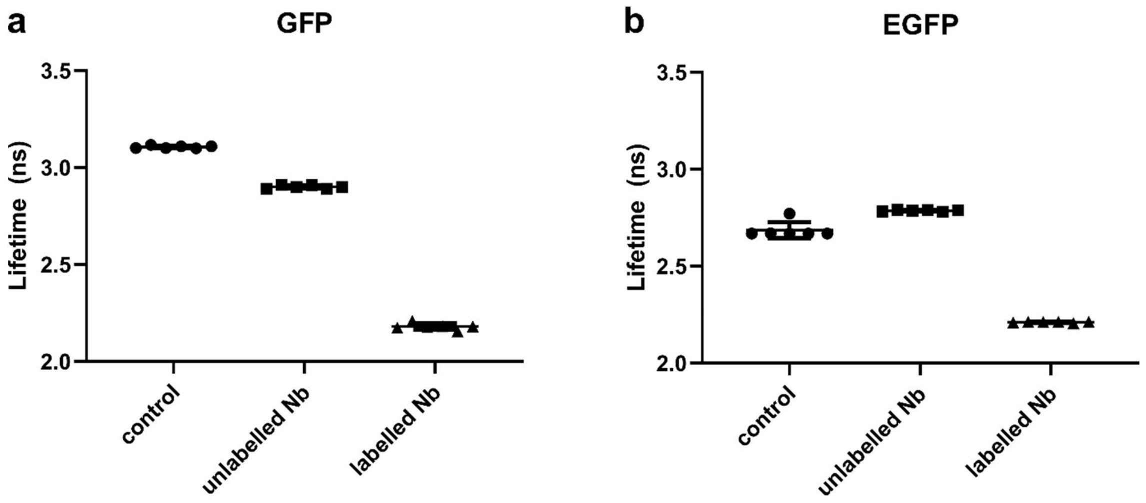


**Supplementary Figure S11: Fluorescence lifetimes in solution.** Fluorescence lifetimes as determined from TCSPC measurements in PBS (pH 7.4) excited with 488 nm for a) GFP and b) EGFP as well as in presence of excess unlabelled or Abberior STAR 635P labelled Nb. Note that for EGFP (control and labelled Nb) a bi-exponentiell fit had to be employed and amplitude weighted lifetimes are given. For all other data a mono-exponential tail-fit was sufficient.

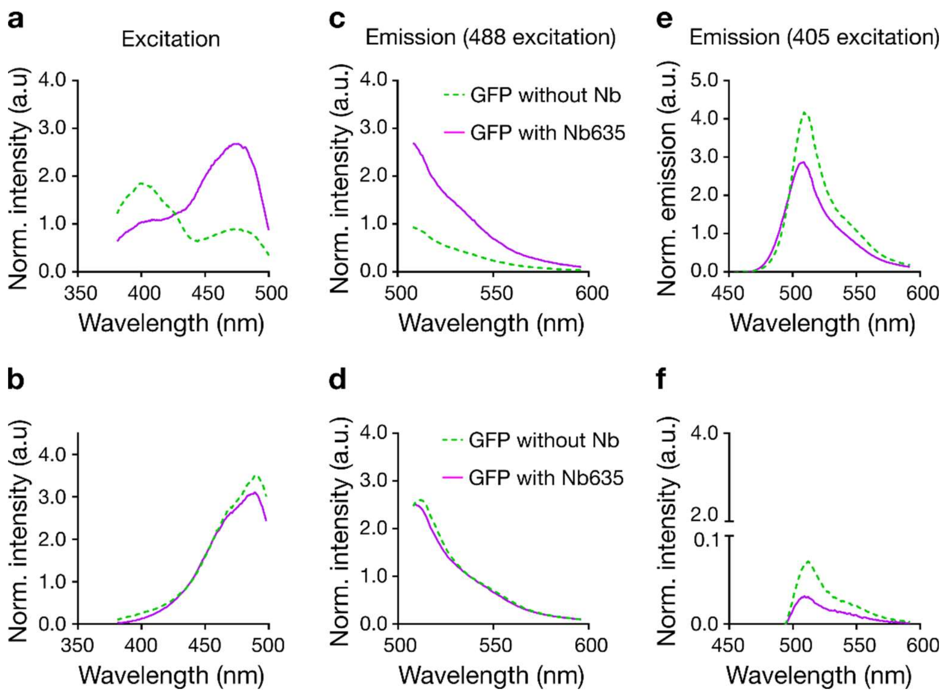


**Supplementary Figure S12: Change in excitation and emission spectra of recombinant GFP and EGFP in solution upon addition of fluorescently labelled Nb.** Excitation spectra for fluorescence detection at 510 - 520 nm (a,b) and emission spectra following 488 nm (c,d) and 405 nm excitation (e, f) of GFP (a,c,e) and EGFP (b,d, f) without Nb (green dashed line) and with labelled AbStar635P-Nb (magenta solid lines). All spectra are averages of three measurements.
